# No fruits without color: Cross-modal priming and EEG reveal different roles for different features across semantic categories

**DOI:** 10.1101/2020.05.22.110338

**Authors:** Georgette Argiris, Raffaella I. Rumiati, Davide Crepaldi

## Abstract

Category-specific impairments witnessed in patients with semantic deficits have broadly dissociated into natural and artificial kinds. However, how the category of food (more specifically, fruits and vegetables) fits into this distinction has been difficult to interpret, given a pattern of deficit that has inconsistently mapped onto either kind, despite its intuitive membership to the natural domain. The present study explores the effects of a manipulation of a visual sensory (i.e., color) or functional (i.e., orientation) feature on the consequential semantic processing of fruits and vegetables (and tools, by comparison), first at the behavioral and then at the neural level. The categorization of natural (i.e., fruits/vegetables) and artificial (i.e., utensils) entities was investigated via cross–modal priming. Reaction time analysis indicated a reduction in priming for color-modified natural entities and orientation-modified artificial entities. Standard event-related potentials (ERP) analysis was performed, in addition to linear classification. For natural entities, a N400 effect at central channel sites was observed for the color-modified condition compared relative to normal and orientation conditions, with this difference confirmed by classification analysis. Conversely, there was no significant difference between conditions for the artificial category in either analysis. These findings provide strong evidence that color is an integral property to the categorization of fruits/vegetables, thus substantiating the claim that feature-based processing guides as a function of semantic category.

## 1. Introduction

The way in which semantic concepts are represented in the brain has been largely informed by neuropsychological studies with brain-damaged patients (for a review, see Mahon & Caramazza, 2009) whose selective impairment in object recognition has been broadly distinguished between natural and artificial (manmade) entities. Notably, however, the category of food—namely, fruits/vegetables—has dissociated from this canonical natural/artificial distinction, with a pattern of deficit that has differentially accompanied an impairment either in the processing of natural entities (i.e., animals; Warrington & Shallice, 1984), artificial entities (i.e., tools; Hillis & Caramazza, 1991) or has demonstrated isolated impairment (Hart, Berndt, & Caramazza, 1985).

Several theories have been proposed to explain the structural organization of concepts in the brain. These theories broadly fall into two general groups. Those that follow a correlated structure principle posit that, while the number of shared versus distinctive features between objects differs across categories, this conceptual distinction is not instantiated at the level of functional neuroanatomy. Those that ascribe to a neural structure principle claim instead that dissociable neural substrates are differentially involved in representing categories (for review, see Mahon & Caramazza, 2009). Proponents of the correlated structure principle assert that the co-occurrence of particular feature types, with an interplay between feature distinctiveness versus sharedness, is what facilitates categorical knowledge and identification (e.g., Conceptual Structure Account; Tyler & Moss, 2001). By extension, categories that possess high within-category similarity, such as that of fruits/vegetables, could be rendered more susceptible to deficit potentially due to a crowding effect of feature overlap that results in low discriminability at the basic level (Humphreys & Forde, 2001).

Alternatively, adherents of the neural structure principle emphasize representational constraints based on the internal neuroanatomical structure of the brain. Some scholars propose a categorical organization of knowledge (Caramazza and Shelton, 1998; Tranel, Damasio, & Damasio, 1997; Damasio, 1990;), claiming that evolutionary pressures imposed functionally dissociable neural circuits dedicated to specific categories (e.g., animals, tools, faces) that have aided our survival. Others, instead, propose that distributed, modality-specific subsystems represent the core organizing principle of semantics; and that these subsystems are differentially important to each category (e.g., Borgo & Shallice, 2003; Warrington & Shallice, 1984). More specifically, they propose that natural entities rely on sensory properties (e.g., color, shape) and artificial entities on functional properties (e.g., use and manipulability) for their classification. Indeed, neurophysiological evidence has revealed an interaction between the processing of category and feature type; lateral portions of the fusiform gyrus have shown to be more active for animals (e.g., natural) as compared to manmade tools (e.g., artificial; Chao, Haxby, & Martin, 1999), with these areas linked to distinct feature processing regions associated with color (i.e., ventral temporal cortex) versus action-related information (i.e., middle temporal cortex; Martin & Chao, 2001), respectively.

Attempts towards resolving the conceptual organization of semantics have primarily focused on a subset of categories, such as animals (i.e., natural) and manmade tools (i.e., artificial), while the category of fruits/vegetables has been relatively under-investigated, despite its relevance to our survival and its inconsistent pattern of deficit that can potentially be mapped to both natural and artificial domains. On the one hand, since fruits/vegetables are (i) ontologically considered to be natural objects and (ii) necessary to our survival, the features important to their recognition should be similar to those of other natural entities, like animals. Indeed, their core semantics seemingly rely heavily on shape and color information (e.g., a banana is elongated, yellow and relatively small, while a watermelon is round, green and relatively large). On the other hand however, fruits/vegetables also frequently engage motoric systems to properly execute eating, which could render them similar to artificial entities and thus relying on function. The current paper aimed to address the role of feature-based processing in the categorization of fruits/vegetables compared to the well-investigated artificial category of tools.

Of the sensory attributes theoretically associated with natural entities, color has garnered particular attention. Some patients have demonstrated dissociation between the perception of color and the knowledge of color typicality associated with objects (Miceli et al., 2001; Samson & Pillon, 2003; Stasenko, Garcea, Dombovy, & Mahon, 2014). A deficit in identifying the appropriate colors associated with fruits/vegetables has either been observed in the presence of intact low-level visual color processing (Samson and Pillon, 2003), or has not disproportionately affected the fruits/vegetables category compared to other living categories (Miceli et al., 2001). However, others have proposed a ‘fractionation’ of visual information relevant within the natural category, with animals relying more on form and fruits/vegetables on color (Breedin, Saffran, & Coslett, 1994; Cree & Mcrae, 2003; Humphreys & Forde, 2001). Indeed, some studies have supported a facilitation of color in the classification and naming of fruits/vegetables (Bramão, Reis, Magnus, & Faísca, 2011; Rossion & Pourtois, 2004), although findings have been equivocal, with some studies either not finding such facilitation (Biederman & Ju, 1988) or showing a greater relative impact for shape information (Scorolli & Borghi, 2015).

Recognition of artificial entities has been shown to rely to a large extent on functional information (e.g., manipulability). Tucker & Ellis (1998) presented participants with an orientation classification task of manipulable objects and demonstrated that objects whose handle was aligned with the responding hand elicited quicker reaction times in orientation judgments (i.e., affordance effect; Gibson, 1977). Such sensitivity to affordance has also been observed in priming tasks in which prior exposure to a line congruent with the graspable axis of a subsequently-presented object has facilitated performance (Chainay, Naouri, & Pavec, 2011). At the neural level, the importance of action-related knowledge to the concept of tools has been argued on the basis of dorsal region activation (associated with the “where” pathway; Goodale & Milner, 1992) to the viewing (Chao & Martin, 2000; Mahon et al., 2007) as well as the naming of tool stimuli (Rumiati, Weiss, Shallice, Ottoboni, & Noth, 2004). Also, the motion associated with tools differentially activates regions of the lateral temporal cortex from that of biological motion (Beauchamp and colleagues, 2002).

In sum, there is ongoing debate as to the neural organization of the semantic system, and how conceptual representations are situated in the perceptual and motoric systems. In this context, there are both experimental results (although not entirely uncontroversial) and theoretical argument to support the hypothesis that visual information (and color, in particular) is critical for natural object representation/processing, whereas motor–related information (e.g., orientation) is critical for manmade object representation/processing. However, the extent to which this asymmetry lies at the core of these concepts is hazy. For example, it is unclear to what extent fruits/vegetables would obey the classic natural vs. artificial dichotomy, and how automatically this asymmetry may arise during semantic processing. These are the issues that we wish to address in this paper.

We report the results of two cross–modal priming experiments in which object images are used to prime lexical decisions on written words. The core manipulation behind our design is that prime images are either (i) normal, (ii) color–modified, or (iii) orientation–modified representations of the relevant objects. Our hypothesis is as follows: if color and orientation are asymmetrically important for natural (fruits/vegetables) and manmade (tools) objects, respectively, then color-modified primes should render a more severe priming reduction for fruits/vegetables, while orientation-modified primes would shrink priming for tools. Alternatively, if fruits/vegetables share overlap with the artificial category, orientation-modified primes should also yield a priming reduction in this category.

Importantly, we presented primes for a short duration (100 ms), with an equally short Stimulus-Onset Asynchrony (SOA; 100 ms), in line with previous work tapping on early semantic processing and the distinction between natural and artificial entities (Dell’Acqua & Grainger, 1999; Kircher, Sass, & Sachs, 2009). Also, the cross modal design was meant to ensure that any generated effect would be genuinely semantic in nature, engaging access across domains.

In Experiment 1, we focused on response times—the experiment was entirely behavioural. In Experiment 2, we collected electrophysiological measures via EEG. A widely used index of semantic processing is the N400 component—a negative deflection in the waveform peaking at about 400ms post stimulus onset, which responds to the semantic ‘predictability’ of a stimulus in a given context (for a review, see Kutas & Federmeier, 2011). This component has commonly been elicited by priming paradigms: greater negativity reflects a reduction in relatedness between primes and targets (Kutas & Federmeier, 2011). N400 effects have been observed for picture-picture priming of real objects (Mcpherson and Holcomb, 1999), and cross-modal priming using both real objects (Kiefer, 2001) and line drawings (Nigam, Hoffman, & Simons, 1992) paired with words. In addition, a graded modulation of the N400 component was reported to reflect a similarly graded modulation of prime-target relatedness (Geukes et al., 2013); this finding is particularly pertinent to the current study, which aims to compare the effect of a prime subjected to visual feature modification on subsequent target processing.

Extending our hypothesis to the EEG domain, we expect that if color is indeed more crucial for natural (fruits/vegetables) entities, color-modified primes should elicit an N400 for this category. Symmetrically, if orientation is critical for manmade entities (tools), then orientation-modified primes should elicit an N400 in this domain. Such findings would suggest that a modification in a feature critical to a given category is perceived as a semantic violation, thus supporting the claim that such a feature is integrally “woven” into the semantic representation of that category. If, instead, fruits and vegetables also critically depend on function (similarly to tools), we may observe a graded N400 negativity, with the degree of negativity modulated by the importance of that feature.

Prior EEG research already speaks to the ability of this technique to distinguish between perceptual and semantic sources of activation (Kiefer, 2001). Moscoso del Prado Martin and colleagues (2006) investigated the spatio-temporal activity patterns of color-related versus form-related words, providing evidence for earlier peak processing of color compared to form. Also, these authors reported that topographies mapped to different underlying neural sources. Furthermore, Amsel and colleagues (2014) utilized a go/no-go semantic decision task to demonstrate that an incongruent attribute of a given object (e.g., purple – lime), when paired together, elicited an N200 component for the no-go condition. This N200 effect was also demonstrated in a similar task for incongruent action-related knowledge referring to graspable objects (Amsel, Urbach, & Kutas, 2013). The authors interpreted these results in support of grounded views of cognition, maintaining that the neural circuitry responsible for perceiving and acting on objects play a role in their conceptual access from long-term memory (Barsalou, 2008).

Furthermore, given the high dimensionality of EEG data, exploratory analysis was performed using linear classification. Previous studies have successfully utilized semantic-decoding algorithms (Simanova, Gerven, Oostenveld, & Hagoort, 2010) to disentangle representations at the neural level.

## 2. Experiment 1

### 2.1. Methods

#### 2.1.1. Participants

A total of 60 healthy right-handed (confirmed by the Edinburgh Handedness Inventory; Oldfield, 1971), native Italian speakers with normal or corrected-to-normal vision participated in the experiment (age range: 18-34 years). Participants were recruited via an advertisement posted on a dedicated social-networking site, and were monetarily compensated for their participation. All participants were naive to the purpose of the experiment and provided informed written consent. The experiment was part of a program that has been approved by the Ethics Committee of SISSA.

#### 2.1.1. Materials

Forty–two Italian words served as critical target stimuli for the experiment. Twenty–one of them represent natural objects (fruits/vegetables; e.g., *pomodoro*, tomato) and twenty–one represent tools (kitchen utensils; e.g., *forchetta*, fork). The main lexical features of these 42 words are illustrated in Table 1a.

**Table 1a.**
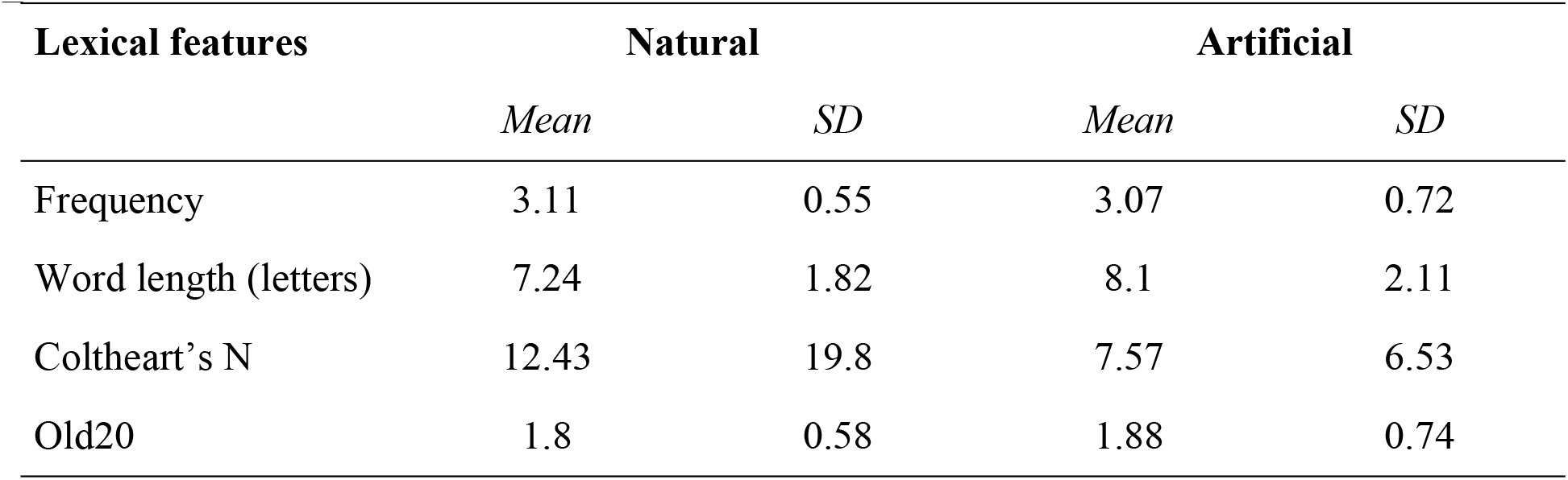
Lexical features for each category. Frequency is given on a zipf scale, which is an improved logarithmic transformation of number of occurrences per million words (van Heuven, Mandera, Keuleers, & Brysbaert, 2014).

| Lexical features | Natural |  | Artificial |  |
| --- | --- | --- | --- | --- |
|  | *Mean* | *SD* | *Mean* | *SD* |
| Frequency | 3.11 | 0.55 | 3.07 | 0.72 |
| Word length (letters) | 7.24 | 1.82 | 8.1 | 2.11 |
| Coltheart's N | 12.43 | 19.8 | 7.57 | 6.53 |
| Old20 | 1.8 | 0.58 | 1.88 | 0.74 |

Each target word was associated with three different prime images, representing the corresponding object: (i) in its canonical color and position (identity prime); (ii) with a clearly non–standard color (color modified prime); (iii) with a clearly non-standard position (orientation–modified prime). In the case of objects without a clear canonical base position (e.g., knife; see (Vannucci & Viggiano, 2000), the original image was chosen to be the one with maximum affordability for a right–hand grasp (Tucker & Ellis, 1998). Identity primes were all taken from the FRIDa database (Foroni, Pergola, Argiris, & Rumiati, 2013).

For the orientation modification, images were rotated 180 degrees in the counterclockwise direction. For the color modification, images were converted to CIELab, which nonlinearly compresses an RGB color image into a three–dimensional coordinate space with the position on one axis representing the red-green opponent channel (a), another the yellow–blue opponent channel (b), and the third the lightness of color, which is the cube root of the relative luminance (Hoffmann, 2003). The advantage of using such a color space is that the lightness contrast can be manipulated independent of a color modification. Furthermore, transformations of images that are high in gray content respond well to manipulations in this color space, meaning that more drastic changes can be achieved as compared to a uni–coordinate modification of hue in the HSL or HSV color space. This was particularly useful for the artificial category, which contained a high number of objects whose principle color was gray. Modifications to both natural and artificial objects were achieved using a custom–written MATLAB^®^ code and made by a fixed proportional change in a– and b–channel value coefficients, the position of each pixel for either axis multiplied by .8 and .2, respectively. This created a purple–blue effect, which we selected among other available options because it rendered a color most naturalistically improbable for our stimuli (e.g., we didn’t have any object among our targets that is prototypically purple or blue). Examples of these prime images can be found in the experimental design schematic of Fig. 1.

**Fig. 1.**
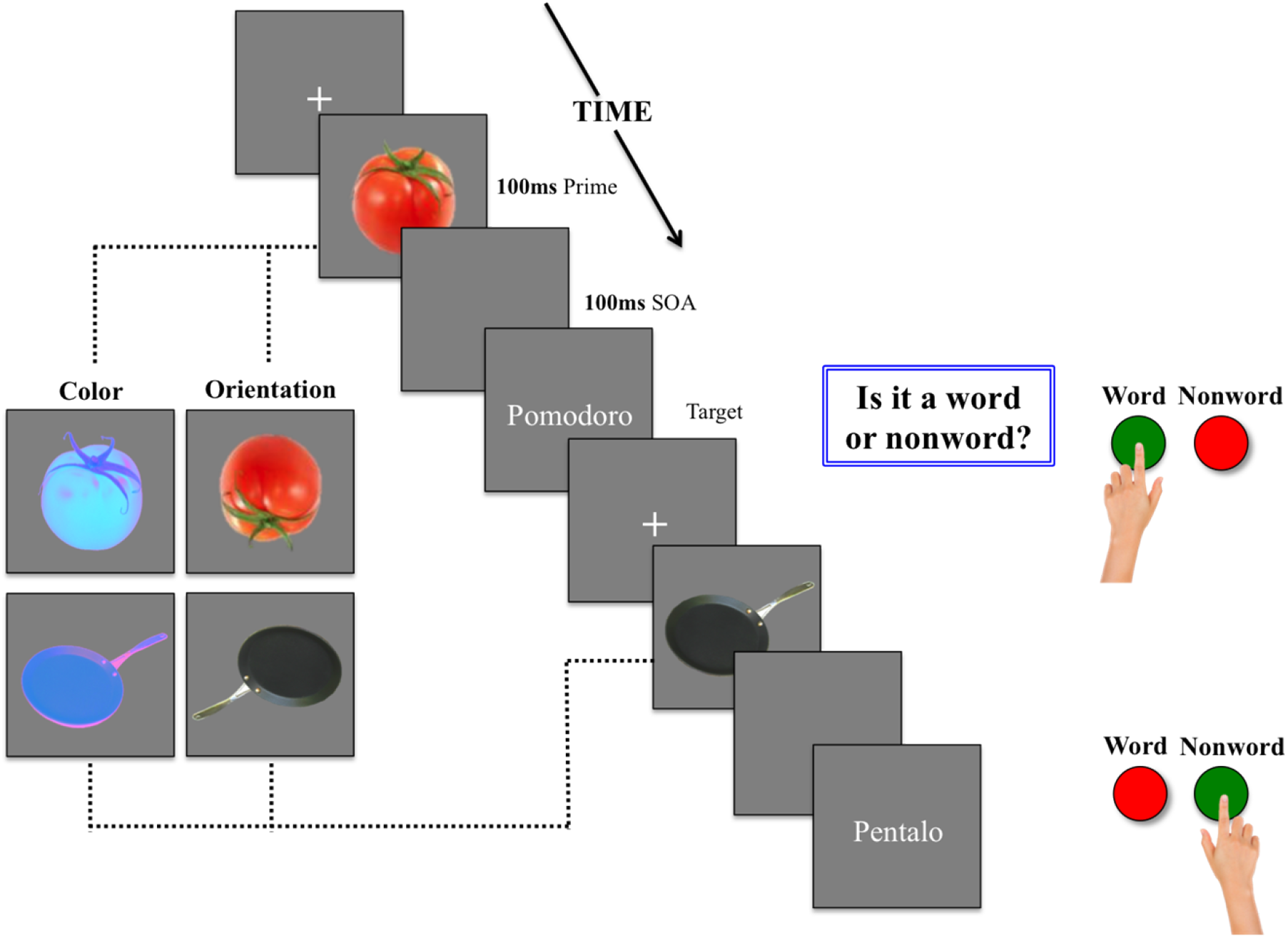
Experimental design schematic of trials of normal condition with color and orientation modification conditions for each category shown to the left. Participants were asked to respond by key press to the question “Is it a word or nonword?” Target strings remained on the screen for 3000ms or until key press.

Identity prime images were matched between natural objects and tools for brightness, spatial frequency, and size in addition to evaluative ratings of discriminability and familiarity, which were based on a 100-point scale. These ratings were obtained from the normative ratings of the FRIDa database. To compute brightness, images were converted to grayscale and the average brightness extracted. To compute object size, objects were isolated using a layer mask in an online image editing application (https://pixlr.com/). Pixels representing the object were converted to black, the background was converted to white, and the ratio between the two was calculated. Spatial frequency was calculated by employing a bi-dimensional fast Fourier transform, which converts the image represented as an array of brightness values to its unique representation in the spatial frequency domain. After transformation, individual pixels represent the power of specific spatial frequencies in a given direction (Foroni et al., 2013). Values for each feature were obtained using custom-written MATLAB^®^ codes (Mathworks, Natick, Massachussets, USA). Both color and orientation–modified images underwent the exact same pipeline (for values of all features, see Table 1b).The assignment of each word target to the three priming conditions was counterbalanced over participants in a Latin Square design, so that all participants received primes from each condition, but saw each target only once.

**Table 1b.**
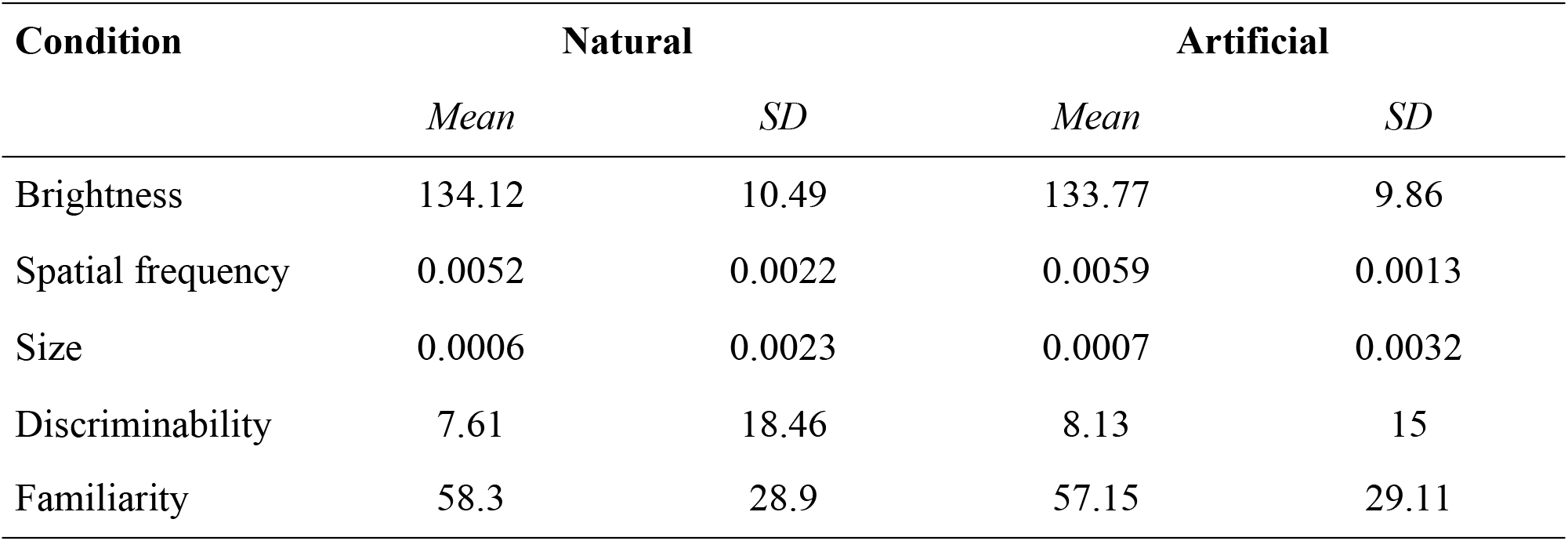
Visual properties and evaluative ratings of stimuli for each category.

For the purpose of the lexical decisions task, 42 nonwords were generated as further target stimuli. They were created from real words by either vowel inversion or single consonant substitution (e.g., *pentalo* from the Italian word *pentola*, pot)—we wanted the task to be challenging, thus engaging participants in deep lexical processing. In order to avoid participants being able to successfully perform the task based on stimuli surface features, words and nonwords were matched for length in letters (Words: *M*=7.67, *SD*=2.02; Nonwords: *M*=7.55, *SD*=1.75) and orthographic neighborhood size (as measured by OLD20; Words: *M*= 1.84, *SD*=0.658; Nonwords: *M*=1.9, *SD*=0.686). These targets were also associated with prime images, similarly to the word–target stimuli. However, nonword-target stimuli were associated with only one image (e.g., normal, color, or orientation) and were not subjected to rotation.

Images were displayed within a 450–pixel square, against a gray background. All letter strings were presented in size 36 Courier New Font, and displayed in white on the same gray background.

#### 2.1.3. Procedure

Participants were tested in a dimly lit room. Participants were seated approximately 60 cm from a monitor with a screen diagonal of 48 cm (resolution: 1280 x 1024 pixels; aspect ration 4:3; refresh rate: 75Hz) and instructed to decide whether or not the letter strings appearing on the screen represented existing Italian words. They were also told that the letter strings would be preceded by a fixation cross, but no mention was made of the presence of the prime images. On each trial, a fixation cross (500ms duration) was followed by the presentation of the prime image (100ms duration), which was followed by a blank screen (100ms duration) and then by the target string of letters. Participants were asked to respond by key press to the question “Is it a word or a nonword?” as quickly and accurately as possible (for a schematic representation including examples of condition manipulations, see Fig. 1). Keys associated with word and nonword trials were counterbalanced across participants. Participants were given six practice trials to familiarize themselves with the task, in addition to the first trial being considered as a practice trial.

Overall, the experiment included 84 trials (42 words and 42 nonwords), for a total duration of about 6 minutes, with the first trial being omitted from the analyses as practice. Stimuli were presented using E-Prime 2.0 Professional (Psychology Software Tools, Sharpsburg, PA).

#### 2.1.4. Analysis

Data were analyzed using mixed–effect modeling as implemented in the *R* package *lme4* (Bates, Maechler, & Dai, 2009).The dependent variable was response time, inverse transformed in order to reduce the typical right skewness shown by RT distributions. We only considered correct responses in these analyses.

Fixed factors were target Category (natural object vs. tools), prime Condition (normal vs. color–modified vs. orientation–modified image) and their interaction. This defines the core structure of the experimental design. *Participants* and *target words* were also specified as random intercept factors, which took into consideration the theoretically uninteresting variation introduced into the data by these factors. This model was confirmed to be the simplest with maximum explained variance by the likelihood ratio test (Pinheiro & Bates, 2000). Robustness to outliers of the model estimates was checked with model criticism as advocated by Baayen, Davidson, and Bates (2008), that is, models were refitted after removing those data points whose standardized residual error was higher than 2.5 and effects were considered significant only if they resisted this procedure). P-values were computed adopting the Satterthwaite approximation for degrees of freedom (Satterthwaite, 1946) as implemented in the lmerTest R package (Kuznetsova, Christensen, & Brockhoff, 2013). In addition, *ggplot2* (Wickham, 2009), *reshape* (Wickham, 2007), and *visreg* (Breheny & Burchett, 2013), were used as part of the R system for statistical computing (Ver. 2.8.1; R Development Core Team, 2013).

### 2.2. Results

Three participants were excluded from the analyses in having exceeded three standard deviations in either reaction time (1 participant) or accuracy (2 participants) performance. Thus, 57 participants (29 female) remained in the final analysis (age: M = 23.67; SD = 3.32). One word was also eliminated from the analysis due to an accuracy of less than 85%. Furthermore, a histogram of individual RTs pooled across all subjects revealed two additional extreme RTs (RT>2500ms), likely to be the result of task inattentiveness, which were thus removed. After removing incorrect trials (n=263; see Supplementary Material Fig. A.1 for accuracies by condition), 4476 data points (2254 word trials) remained in the final analysis. The overall accuracy rate for the task was 94.45%. Means, medians and standard deviations for response times in the six conditions are illustrated in Table 2a.

**Table 2a.** Raw RT means, medians, and standard deviations for Condition x Category.

| Condition | Natural |  |  | Artificial |  |  |
| --- | --- | --- | --- | --- | --- | --- |
|  | *Mean* | *Median* | *SD* | *Mean* | *Median* | *SD* |
| Normal | 672.67 | 614 | 221.06 | 759.79 | 702.5 | 258.83 |
| Color | 705.41 | 646 | 230.66 | 779.24 | 719.5 | 276.92 |
| Orientation | 684.98 | 625 | 235.51 | 804.28 | 735.5 | 292.36 |

Statistical modeling revealed a significant effect of target Category [F(1, 38.93) = 7.7, p = .008], prime Condition [F(2, 2842.39) = 6.86, p = .001], and, most importantly, their interaction [F(2, 2842.31) = 4.47, p = .01]. To explore the interaction, we fitted separate models for natural and artificial target words. For the natural category, color-modified primes generated a significant reduction in priming compared to orientation–modified (t_(56)_ = −2.12, p = .034) and normal primes (t_(56)_ = −3.87, p < .001), which in turn didn’t differ significantly from each other (P = .084). For the artificial items instead, orientation-modified primes penalized priming the most, providing less priming than both color–modified (t_(56)_ = −2, p = .045) and normal primes (t_(56)_ = −2.88, p = .005), which in turn didn’t differ from each other (P = .38).

Reaction times were also binned into quartiles to explore the effect across the distribution. F-statistics and p-values for the effect of condition across quartiles, divided by category, is provided in Table 2b. An interesting finding is that, for the natural category, a visualization of the data revealed a similar pattern of performance across all response time intervals. However, for the artificial category, the main effect of condition emerged at later time intervals, representing a distributional shift. Such a shift in RT distribution could potentially be attributed to retrospective retrieval processes, where the target is semantically matched to the prime (Yap, Balota, & Tan, 2013).

**Table 2b.** F-statistics for anova of models divided by category and quartile.

| Quartile | Natural | Artificial |
| --- | --- | --- |

|  | *F-value* | *Pr(>F)* | *F-value* | *Pr(>F)* |
| --- | --- | --- | --- | --- |
| Q1 | 13.52 | <.001 | 1.59 | .206 |
| Q2 | 15.46 | <.001 | 4.42 | .02 |
| Q3 | 14.87 | <.001 | 12.7 | <.001 |
| Q4 | 1.45 | .235 | 5.69 | .004 |

### 2.3. Discussion

We used a cross–modal, lexical decision priming task to test the hypothesis that natural and artificial entities rely on different properties for their semantic processing. We have shown that, indeed, the processing of natural entities is more sensitive to modifications of a sensory property (i.e., color), while that of artificial entities is more sensitive to modifications of a functional property (i.e., orientation).

The cross-modality of the effect (primes were object images, while targets were words) further suggests that the effect described here is semantic in nature. Moreover, both words and nonword targets were constructed from real words representing the prime object by a single vowel/consonant change; in this way, both word and nonword targets were semantically related to the prime image. Neely (1977) described a retrospective matching strategy in which checking whether a target is related to a prime may induce priming in the lexical decision task. As nonword targets are typically unrelated to the prime, the prime-target relatedness proportion is lower for nonwords. as the thus reflecting the same prime-target relatedness proportion for each. In the current study, the relatedness proportion between prime and target was 1 for both words and nonwords, meaning that participants could not have employed a retrospective matching strategy to identify words differentially from nonwords based on a semantic relationship that only target words share with the prime (de Wit & Kinoshita, 2015). This is generally considered as further confirmation that priming derives from access at the conceptual level.

While color is most integral to the processing of fruits/vegetables and orientation to the processing of tools, there were also reductions in priming for each of the modified conditions with respect to the normal condition. This suggests that, while a certain feature may be of critical importance to a particular category, other features may also play a role in its conceptual representation. Furthermore, this reduction was more evident for fruits/vegetables, which displayed greater susceptibility to priming interference by an orientation transformation than utensils did to a color change. This is not particularly surprising after all; as hypothesized, fruits/vegetables can also be considered graspable entities that invoke motor affordances (although such an effect has been reported to depend on overt responses to graspability, which were not involved in this task; Netelenbos and Gonzalez, 2015).

## 3. Experiment 2

In Experiment 2, we utilized EEG to test whether the interaction between category and feature type would manifest at the neural level, as indexed by the N400 component. We hypothesized that a modification of a feature integral to the representation of a given category would be processed as a semantic violation, thus eliciting an increase in negativity in the N400 time window.

### 3.1. Methods

#### 3.1.1. Participants

Forty native speakers of Italian (32 after exclusion; see *Data* section below) partook in the experiment. Participants were healthy, right-handed (confirmed by the Edinburgh Handedness Inventory; Oldfield, 1971) individuals (mean age ± standard deviation= 24.33 ± 2.45; range= 19-30 years; 17 females/15 males), recruited via an advertisement posted on a dedicated social-networking site and monetarily compensated (25€) for their participation. All participants had normal or corrected-to-normal visual acuity, no history of neurological or psychiatric illness, and no history of drug or alcohol abuse that might compromise cognitive functioning. All participants were naive to the purpose of the experiment and provided informed written consent. The experiment was part of a program that was approved by the ethics committee of SISSA.

#### 3.1.2. Materials

The materials used in Experiment 2 were identical to those employed in Experiment 1.

#### 3.1.3. Methods

Participants were seated approximately 60 cm from a monitor with a screen diagonal of 48 cm (resolution: 1280 x 1024 pixels; aspect ration 4:3; refresh rate: 120Hz). Participants were presented with a lexical decision-priming paradigm identical to the one used in Experiment 1 with one exception-rather than indicating their response by button press, they were asked to silently categorize the string of letters as word or nonword. However, on approximately 25% of trials, a question mark appeared following the target string, prompting a vocalized response. This was done to maintain task engagement while minimizing any motor-related contamination introduced by a button press (Vliet et al., 2014). Each target string was associated with one question mark to ensure that participants could correctly classify all target strings.

Overall, one block of the experiment consisted of 84 trials (42 words and 42 nonwords), identical to Experiment 1, and was repeated five times. Stimuli were presented using E-Prime 2.0 Professional (Schneider, Eschman, & Zuccolotto, 2012). The experiment was performed in a sound-proof cabin. All electrical devices that were not sources of direct current and could interfere with EEG wave acquisition were turned off before experiment onset.

#### 3.1.4. EEG Acquisition

Continuous EEG was recorded from an array of 128 silver-chloride Biosemi active electrodes mounted on an elastic cap (topographic placement: radial ABC layout system). Two external electrodes were placed on the left and right mastoids (A1, A2) as reference. However, due to high impedance values and signal noise discovered upon manual inspection, mastoids were later discarded and average reference used (i.e., mean of all electrodes). EEG signal was amplified using a Biosemi Active-Two amplifier system (Biosemi, Amsterdam, Netherlands) at a sampling rate of 1024 Hz. An electrode located near Cz (common mode sense: CMS) was used as the recording reference. The direct current offset was kept below 25 mV. Data acquisition was made using the software Actiview605-Lores (www.biosemi.com).

#### 3.1.5. EEG Preprocessing

EEGLAB (Delorme & Makeig, 2004) and Fieldtrip (http://www.ru.nl/neuroimaging/fieldtrip)-open source Matlab toolboxes-were used to perform all preprocessing steps. Off-line data preprocessing included a digital high-pass filter of 0.1 Hz (as recommended by Tanner, Morgan-Short, & Luck, 2015) and a low-pass filter of 100 Hz. Data were down–sampled to 256 Hz and segmented into epochs of 1200 ms, starting 200 ms before prime onset. Before proceeding with data cleaning, incorrect trials were identified and removed. Incorrect classification of a target string resulted in all trials in which that string appeared being removed. Electrodes exceeding a certain z-score threshold (kurtosis = 4; probability = 4; spectrum = 3) were removed from the data (Delorme, Sejnowski, & Makeig, 2007). A subset of electrodes sensitive to ocular movements were withheld from this threshold check and later passed to Independent Component Analysis (ICA). Noisy trials were deleted by visual inspection and data were referenced to the average. ICA using the Infomax algorithm was performed on all data for elimination of artifacts related to ocular and muscular movements. Data were time-locked to the onset of the prime image and baseline (200 ms pre-prime onset) removed. Automatic trial rejection was then performed for a more fine-grained cleaning: ±50 dB threshold in the 0-2 Hz frequency range (for capturing residual eye movements) and +25 to −100 dB in the 20-40 Hz (for capturing muscle movements). Finally, missing electrode data were interpolated to the original 128-electrode montage.

#### 3.1.6. Data

Of the initial 40 participants, 32 were retained for the analysis. One participant was eliminated due to high error rate (8.3% trials) and the remaining seven due to excessive noise (> 25% of trials were rejected). Error rate was less than 1% (*M* = 0.7; *SD* = 1.2) and the average number of trials rejected due to noise was less than 15% (*M* = 86.67%; *SD* = 6.6%) for the remaining subjects.

#### 3.1.7. Standard ERP Analysis

ERPs were computed for epochs extending from 200 ms pre-prime onset to 1000 ms post-target onset. Analysis was performed using threshold-free cluster-enhancement (TFCE), which is a cluster-based technique that embeds permutation-based statistics for significance testing (Mensen & Bern, 2014). The advantage of TFCE over other cluster size/mass techniques is that it uses information about both the intensity and the spatial distribution of the signal to enhance weak, but broadly supported signals to the same numerical values as strong, but highly focal signals, without having to select a cluster-forming threshold a priori (for a more thorough description, see Mensen & Bern, 2014). In addition, this technique outperforms other clustering methods in controlling for Type 1 errors (Pernet, Latinus, Nichols, & Rousselet, 2015). The entire epoch was submitted to TFCE analysis, which resulted in a significance matrix (electrode x time) in which clusters were identified based on the span of significance in both time and space. The cluster of interest was the N400 time window, which was pre-defined as a window spanning from 250-500ms post-target onset (Kutas & Federmeier, 2009).

The first comparison of interest was between all word and nonword trials, irrespective of prime pairing. This was to ensure that our paradigm indeed elicited the most basic semantic distinction between meaningless and meaningful stimuli as previously demonstrated in the literature (Kutas & Federmeier, 2000). Average waveforms for all word and nonword trials were calculated for each participant and a grand average computed to compare conditions. Thereafter, nonword trials were discarded from further analysis and only word trials considered. Word trials were divided by category into those representing natural versus artificial entities. For each category, all combinations of conditions were compared (i.e., normal–color, normal–orientation, color–orientation).

#### 3.1.8. Classification

In addition, we performed a searchlight classification on all electrodes in the interval of 250 to 500 ms post-target onset, corresponding to the N400 time window (Kutas & Federmeier, 2009). We chose linear discriminant analysis (LDA) classification because it employs an algorithm that is able to handle input of more than two classes; LDA utilizes a data reduction technique to divide the feature space based on maximum variance, similar to Principle Component Analysis (PCA), and generates weights to discriminate between classes (Subasi & Gursoy, 2010). LDA classification was performed separately for the natural category (all conditions included) and then artificial category (all conditions included).

Classification was performed at the individual participant level. For each participant, individual trials, serving as *observations*, were submitted to the classifier. In cases where conditions contained different numbers of trials, a random subset of trials from the larger condition was sampled to match. Algorithms employed a cross-validation measure and an odd-even partition scheme whereby both even and odd runs served as training and testing sets. A searchlight approach was used to create neighborhoods of channels with each channel serving as ‘origin’ and identifying neighboring sensors within a certain configuration. Neighbors were defined based on a triangulation algorithm in Fieldtrip that allows for the building of triangles of ‘nodes’ that are independent of channel distances. Classification was then performed on each searchlight neighborhood across the entire scalp. Accuracies were generated per comparison, per participant, which were then thresholded for significance at the group level using Monte Carlo permutation testing with 10000 iterations. Essentially, permutation testing in this context involves subtracting the chance accuracy from the computed accuracies and randomly flipping the sign over iterations. This generates multiple t-statistics over iterations that are then corrected for multiple comparisons using TFCE, which results in a z-score map that can be plotted for significance. Significance was defined as a one-tail probability value of z > 1.64 (p < 0.05).

### 3.2. Results

#### 3.2.1. ERP results

In a first comparison between word and nonword trials, TFCE analysis showed that nonword trials generated greater negativity relative to word trials for 19 unique channels in the left fronto-central region for the time range of 288 – 423 ms (peak channel at 389 ms: C1; T = −6.27; p < 0.002). This finding is in line with previous semantic literature that has demonstrated an N400 effect for meaningless compared to meaningful stimuli (Kutas & Federmeier, 2000).

Examining word trials only, for the natural category, TFCE analysis revealed a significant difference between the color and normal condition, such that color trials elicited significantly greater negativity than normal trials for 11 unique channels in the left fronto-central region in the time range of 213 – 387 ms (peak channel at 256 ms: C1; T = 6.12; p < 0.009). The resulting brain topography and grand mean waveforms for all three conditions can be seen in Fig 2. Next, although an evident difference emerged in the grand mean for selected electrodes between the color and orientation conditions as well, TFCE confirmed that the color condition elicited a greater N400 response with respect to the orientation condition. This effect was found for 8 unique channels in the central region for the time range of 299 – 381 ms (peak channel at 320 ms: B1; T = −6.07; p < 0.001; see Fig. 3). These findings are in line with our hypothesis that a color-modification for fruits/vegetables should yield a semantic violation as indexed by an increase in N400 amplitude. Last, a comparison between orientation and normal conditions revealed one channel that exhibited significantly higher amplitude compared to the normal condition, in the centro-parietal region for the time range 89 – 97 ms (peak channel at 93 ms: Pz; T = −5.88; p < 0.05).

**Fig 2.**
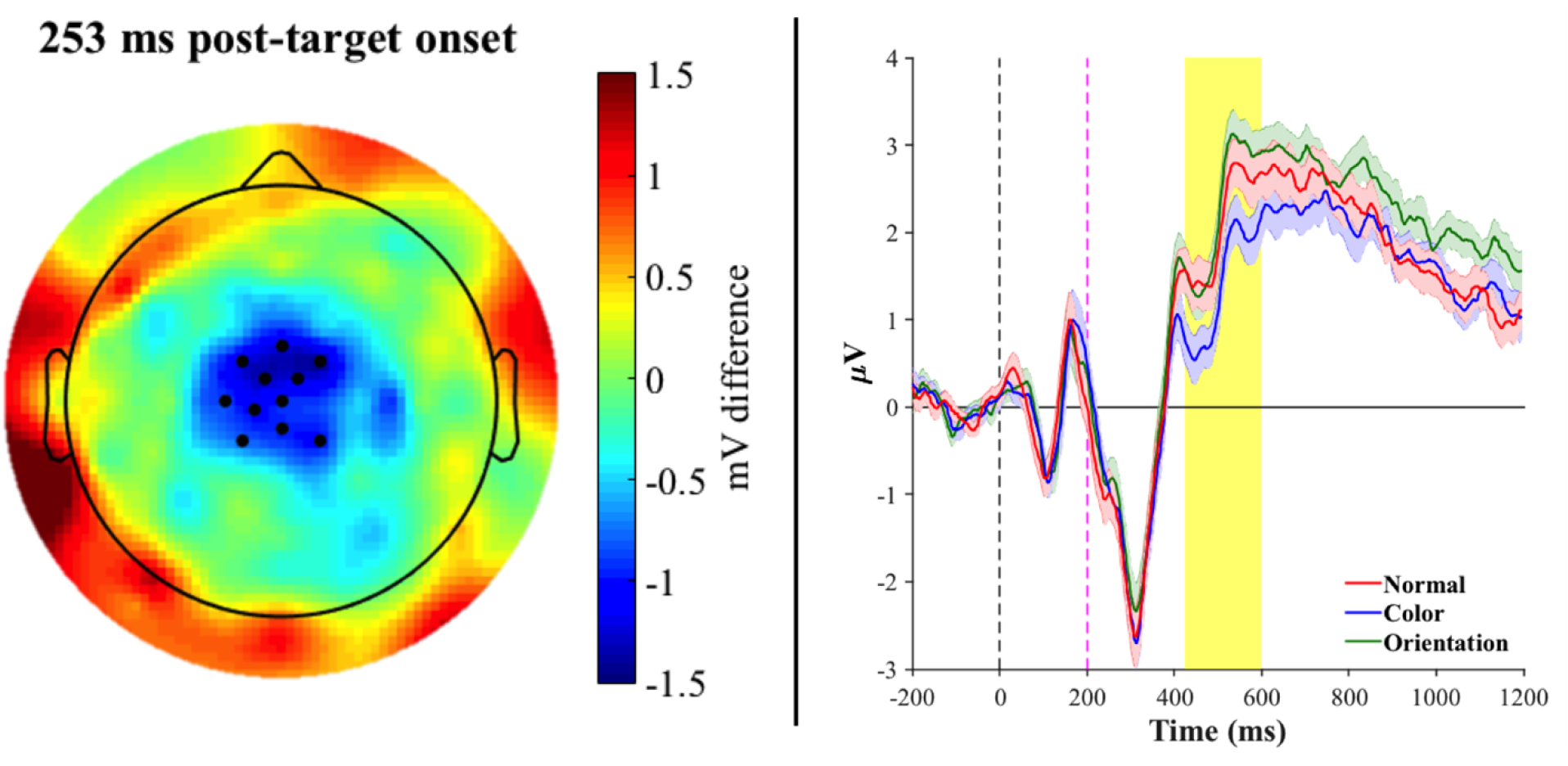
Left panel: Topographic map of the brain at peak channel significance (~253 ms post-target onset). Topography reflects the difference between the grand average for Color – Normal trials, black dots represent the 11 unique significant electrodes at p < 0.009. Right panel: Mean waveforms across the 11-electrode cluster plotted separately for the normal (red), color (blue), and orientation (green) conditions. Significance reflects the difference between the color and normal condition (orientation is also plotted for visualization of the pattern of effect). Shaded regions along the waveforms indicate standard error of the mean. The shaded yellow bar denotes the N400 time window. The orientation condition elicited a greater positivity in the N400 time window with respect to the normal condition, whereas the color condition elicited the canonical N400.

**Fig 3.**
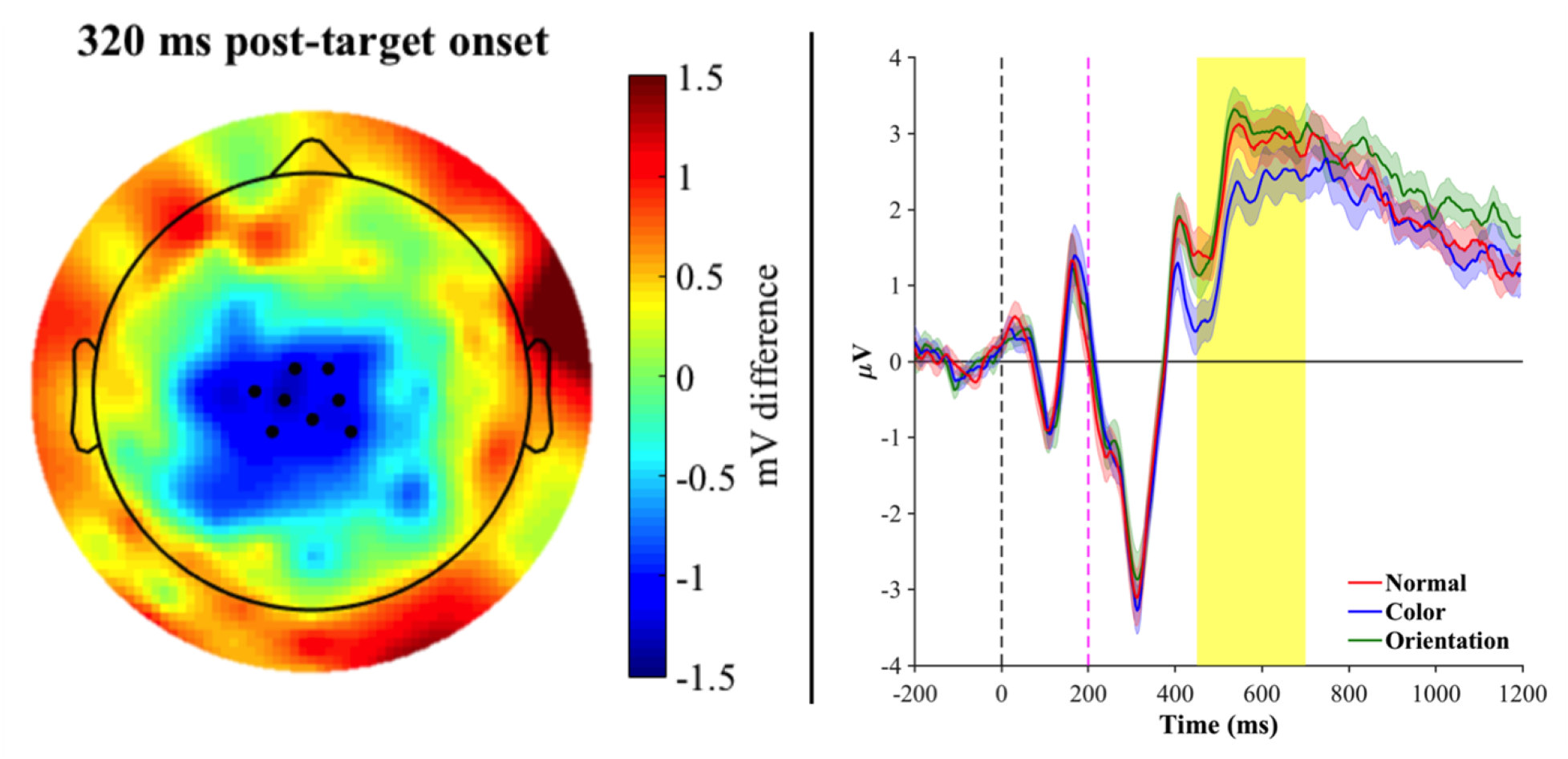
Left panel: Topographic map of the brain at peak channel significance (~320 ms post-target onset). Topography reflects the difference between the grand average for Color – Orientation trials, black dots represent the 8 unique significant electrodes at p < 0.001. Right panel: Mean waveforms across the 8-electrode cluster plotted separately for the normal (red), color (blue), and orientation (green) conditions. Significance reflects the difference between the color and orientation condition (normal is also plotted for visualization of the pattern of effect). Shaded regions along the waveforms indicate standard error of the mean. The shaded yellow bar denotes the N400 time window. The color condition, compared to the orientation condition, elicited the canonical N400.

For the artificial category, the color compared to normal condition revealed a significant cluster of 4 unique channels displaying greater negativity for the color compared to normal condition (peak channel at 102 ms (post-prime): A26; T = −6.6; p < 0.03). This effect was of short duration, spanning 8 milliseconds, and only witnessed in the prime window. A comparison between the orientation and normal condition did not reveal significant channels at any time point (P > .1), nor did the comparison between the orientation and color condition.

#### 3.2.1. ERP results

#### 3.2.2. Classification results

A statistical z-score map of LDA accuracies can be found in Fig. 4. For the natural category, the LDA classifier successfully discriminated between the three conditions in 7 channels as origin in the searchlight (z-range: 1.7–2.18; p-range: .015–.045). These channels were located in the left fronto-central region and resembled the ERP scalp topography of significant channels for the word-nonword comparison as well as the natural category comparison, albeit to a lesser extent. These results confirm the robustness of the main finding that an exposure to a prime manipulation in a feature relevant to the categorization of the natural category of fruits/vegetables yields an effect in the N400 time window.

**Fig. 4.**
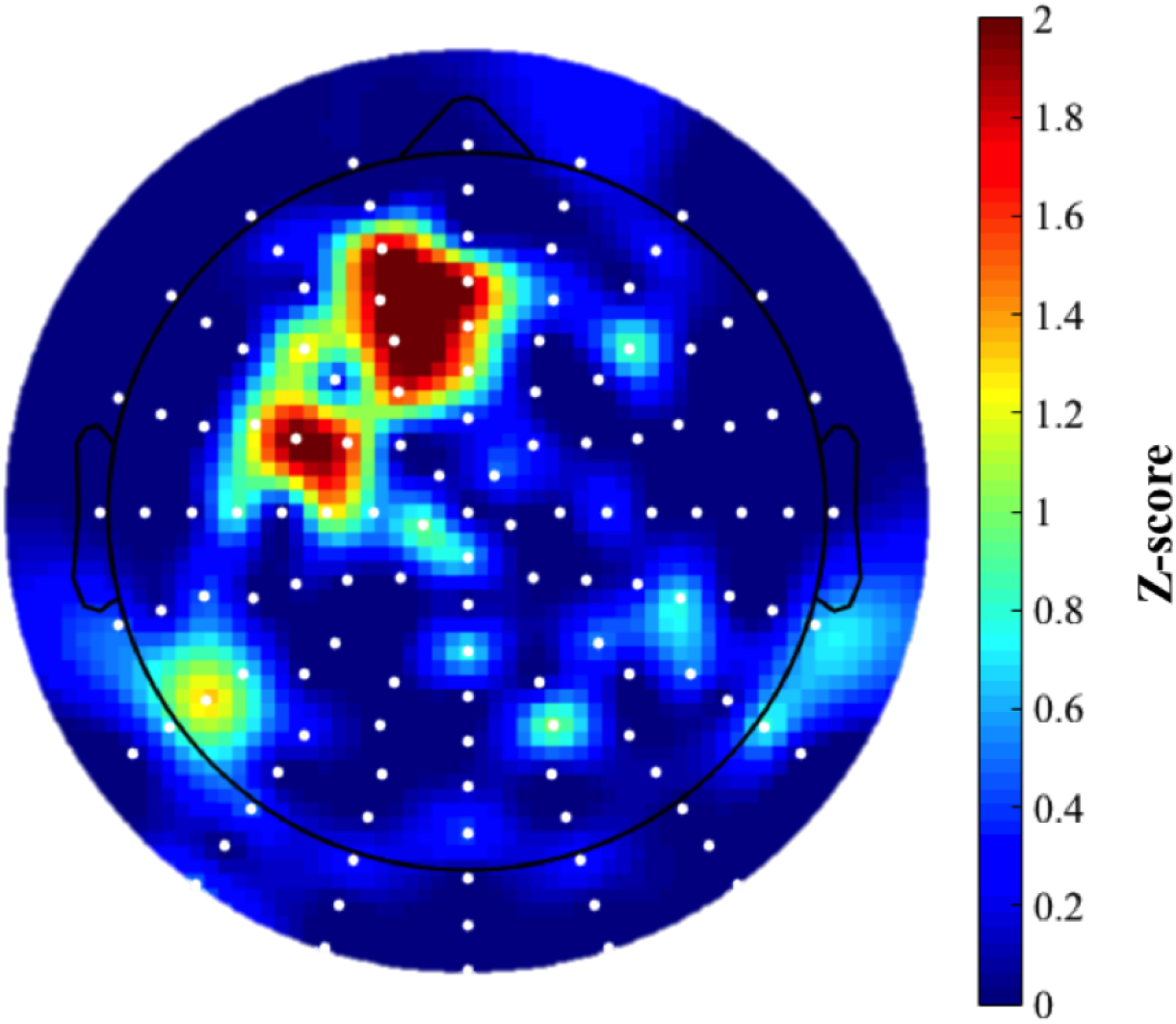
A statistical z-score map of LDA accuracies for the natural category. Significance was defined as a one-tail probability value of z > 1.64 (p < 0.05) and can be observed in 7 channels that served as searchlight origins (dark red regions) in left fronto-central sites.

For the artificial category, although there was no significant difference between conditions when comparing ERP amplitudes, one channel neighborhood was able to significantly discriminate between classes (z = 1.9, p = .029).

## 4. Discussion

ERP component analysis demonstrated a significant effect in the N400 time window (i.e., 250 – 500 ms post-target onset) that was modulated by the manipulation in prime for fruits/vegetables. A comparison of the difference wave [orientation – normal vs. color – normal] revealed a significant negative deflection for the color-modified condition, in line with our predictions and with the results of Experiment 1. Interestingly, it was observed that the magnitude of this difference was driven by a positive deflection of the orientation–modified condition with respect to the normal condition. Given what appeared to be a trend towards a graded nature of a priming effect observed in the behavioral data of Experiment 1, it was hypothesized that an orientation-modification may also elicit an N400 effect, albeit to a significantly lesser extent than color. However, we did not observe such N400 modulation. One likely explanation is that although functional knowledge could still be relevant to the concept of fruits/vegetables, it may have a neural signature that cannot be captured by the N400 component alone (e.g., propagation from an earlier component; Mcpherson and Holcomb, 1999).

Quite interestingly, when we employed a multivariate data analysis tool, the LDA classifier was indeed able to accurately distinguish between the three conditions at fronto-central channel sites with a slightly left-lateralized bias; this pattern of results resembled the ERP scalp topography of significant channels for the word-nonword comparison as well as that of the color vs. normal condition comparison. LDA’s ability to discriminate between conditions could be due to the fact that, similarly to PCA, it projects the data into a space that is based on feature extraction of dimensions common to all input, to avoid redundancy and maximizes the difference between classes. However, as this analysis utilizes information at the individual trial level, it may be more sensitive to capturing differences between conditions that do not significantly emerge at the mean level.

Conversely, ERP component analysis of the artificial category did not reveal a significant difference in the time window of interest. The only finding was a slight shift and reduction in the P100 (P1) component to the color compared to normal condition, which did not survive correction for multiple comparisons. In any case, this reduction could easily be due to low-level perceptual differences in the prime image as the P1 component has shown to be sensitive to external properties of the stimulus like luminance, which can consequently exert an influence on attention (Luck, Woodman, & Vogel, 2000). The fact that there was no observable N400 could be due to a number of reasons. One possibility is that action knowledge relating to manipulability may not be automatically engaged at the conceptual level. Indeed, a recent study by Yu and colleagues (2014) challenged the automaticity of an affordance effect previously reported by Tucker and Ellis, replicating their results only when explicit task instructions were given to imagine interacting with the object. This suggests that relevant motor regions may be activated only under specific circumstances such as goal-directed behavior. However, given the pattern of behavioral results that would oppose this claim, it may be more plausible that factors interfering with capturing the neural response are, in fact, responsible. For instance, higher values of Coltheart’s N, which is a measure of orthographic neighborhood size (Coltheart, Davelaar, Jonasson, & Besner, 1977), has been shown to elicit N400s of smaller amplitude (Holcomb, Grainger, & O’Rourke, 2002). The artificial category did possess higher values of Coltheart’s N with respect to the natural category, which could have mitigated an effect. Given the several other constraints that we had on our stimuli selection (e.g., frequency and length matching, items needed to be easily picturable and have a characteristic color), it was impossible to work out a better matching on this front.

LDA classification was able to significantly discriminate between conditions in the N400 time window at one right posterior channel site. Interestingly, a recent study by Hauser and colleages (2015) combined fMRI with EEG to show the scalp topography related to the hand motor area as defined by ROI analysis and found a similar pattern of activity at right-lateralized posterior sites. The fact that an effect did not emerge in the ERP amplitudes could most likely attest to the loss of information when trials are averaged across conditions, necessitating a finer-grained (multivariate, across time and space) analysis (Murphy, Poesio, Bovolo, Bruzzone, & Dalponte, 2011). This finding suggests that information in the prime window did exert a differential effect on the N400 time window, such that meaningful information distinguishing between conditions was contained at this channel site despite the absence of an effect at the level of grand mean. However, that only one site could accurately distinguish between classes could also be the result of color not being as relevant to the artificial category; while some utensils may also possess a degree of color diagnosticity, there may not have been enough coherence within the category to allow for a robust decision boundary to be drawn by the classifier.

## 5. General Discussion

The objective of the current study was to assess if feature-based properties of fruits/vegetables and tools were integrated with object knowledge at the conceptual level. To this end, we administered a lexical semantic priming paradigm and recorded reaction times and neural responses to investigate the processing of an object when a modification of a feature potentially crucial to its conceptual representation has been presented. Analysis of the reaction time data of Experiment 1 supported the claim that the presentation of sensory or functionally relevant features can differentially enhance or detract from the priming effect depending on the type of category. Such evidence are in line with the sensory-functional hypothesis as first proposed by Warrington and Shallice (1984), whereby (i) concepts are *grounded* in the sensory and functional subsystems that are relevant to their processing and (ii) these subsystems are differentially important depending on category type. Moreover, the neural data, particularly for fruits/vegetables (FV), corroborates this claim as both standard ERP analysis and multivariate techniques revealed a significant modulation of semantic processing by color-modified primes.

Fruits/vegetables have proven to be a very interesting, yet still controversial category in the semantic literature. While ontologically considered to be a member of natural kinds, neuropsychological patient studies, in addition to work with healthy individuals, have shown it to be a category that, in some respects, traverses a coarse natural/artificial domain distinction (for a review, see Capitani, Laiacona, Mahon, & Caramazza, 2003). In a review of posterior cerebral artery patients, Capitani and colleagues (2009) suggested that the underlying neural substrate for the processing of fruits/vegetables may be different than that of animals; specifically, findings from these patients suggest that middle fusiform lesions disproportionately impair plant life (including fruits/vegetables), whereas anterior temporal lesions that are typically the result of herpes simplex encephalitis create a disproportionate deficit for animals. A fractionation between animals and fruits/vegetables has also been proposed at the level of visual information, with the former relying more on form and the latter on color for semantic processing (Breedin et al., 1994; Cree & Mcrae, 2003; Humphreys & Forde, 2001). While studies have supported a facilitation by color of the processing of fruits/vegetables (Bramão et al., 2011; Rossion & Pourtois, 2004), results have been equivocal, with studies either not reporting such facilitation (Biederman & Ju, 1988) or showing a greater impact of other modal information to its processing (Scorolli & Borghi, 2015). The current study demonstrated that, indeed, color is an integral feature to the representation of fruits/vegetables at the conceptual level—response times during lexical-semantic processing of fruits/vegetables were primed substantially less when a target word was preceded by color-modified primes of the same object, and electrophysiology nicely backed up this behavioral pattern of results.

An interesting, though not fully unexpected, finding was that LDA classification of the waveforms could accurately distinguish between all three conditions of fruits/vegetables, even if ERP (or response times) didn’t reveal significantly strong difference between orientation-modified and normal pictures. This suggests that orientation may also be relevant to the categorization of fruits/vegetables, possibly because these are graspable entities with which we must interact for our survival. The fact that a significant difference only emerged at the level of classification could be due to the fact that only some fruits/vegetables necessitate an appropriate affordance grasp for their consumption (e.g., a banana may necessitate more manipulation than a grape). It has been shown that fruits/vegetables can also invoke motor affordances in the categorization process, although such an effect has been observed for overt responses to graspability (Netelenbos and Gonzalez, 2015). It has, however, been suggested by Gainotti and colleagues (2010) that the left lateralization of lesions underlying a deficit in the processing of fruits/vegetables in posterior cerebral artery patient profiles (see Capitani et al., 2009) may reflect the reliance on motor knowledge that is necessary for eating actions. Here, we demonstrate that the mere viewing of fruits/vegetables may automatically-activate motor affordance. Although orientation appears relevant to the semantic processing of fruits/vegetables, it is so to a lesser extent than color. This finding lends further support for a sensory-functional theory of semantics over, for instance, a domain-specific approach that should theoretically predict a similar performance across modality as the organizing principle of semantics is assumed to be at the level of category, rather than modality.

As for the artificial category, reaction time data revealed a significant slowing of lexical-semantic processing when words representing tools were preceded by orientation-modified primes. This finding further confirms the differential importance of modality-specific information to the conceptual representation of an object that necessitates that modality for their processing (Bonner & Grossman, 2012). At the neural level, although a modulation by feature type did not manifest in the standard ERP analysis, LDA classification did reveal a difference in one right-lateralized posterior channel site. Apart from the fact that averaging across conditions could have led to the abolishing of an effect for which a finer-grained analysis may be necessary (Murphy et al., 2011), it could be that greater variability within the category of tools rendered some exemplars more reliant on motor affordances than others (Chen et al., 2016). In a similar vein, potential variability in color diagnosticity, albeit less important to the artificial category, could have been responsible for LDA performance in distinguishing between conditions. While this greater variability could have driven successful LDA classification, but not significant effects in component analysis, on the flip side, it could also be the reason why only one channel site could successfully perform this distinction. In one way, this finding does complement the behavioral pattern where, although RTs demonstrated a significant difference between normal and orientation conditions, there were also priming reductions observed for the color condition.

## 6. Conclusion

In the current study, response times, ERP waves and multivariate techniques applied to the EEG signal converged to indicate a direct involvement of perceptual color processing and, to a lesser degree, orientation processing on conceptual access to fruits/vegetables.

Although the inverse pattern of reaction time emerged for the artificial category, this did not manifest in the neural pattern of N400; however, a more fine-grained analysis of the electrophysiological data could accurately distinguish between conditions. Our results, in sum, support modal theories, whereby modality-relevant input is integrated at the conceptual level. As the present study only considered two features that have been prominent in the semantics literature, future studies should consider the role of other modalities (e.g., taste, texture) that may be intrinsic to the conceptual representation of natural and artificial objects.

## 7. Acknowledgements

We would like to thank Arash Fassihi, Yamil Vidal Dos Santos, Romain Brasselet, and Sebastian Korb for their support at various stages of the scripting process. We would also like to again thank Arash Fassihi and Romain Brasselet for their insight into the analytic approach.

## 10. Supplementary Material Captions

Appendix A. Supplementary Data:

Fig. A.1. Violin plot of accuracies averaged across all trials per subject, plotted by condition. Each dot represents the mean reaction time for an individual subject. Solid purple line denotes the mean of each condition.

